# Prefrontal Control of Innate Escape Behavior – A Neural Mechanism of Enhanced Posttraumatic Threat Detection

**DOI:** 10.1101/2022.12.21.521361

**Authors:** Ami Ritter, Shlomi Habusha, Shahaf Edut, Oded Klavir

## Abstract

Innate defensive responses, while primarily instinctive, must also be flexible and highly adaptive to changes in risk assessment. As such, efficient innate escape behavior requires intricate processing to minimize reaction time while maximizing the success and adaptivity of the action. The superior colliculus (SC) is a subcortical sensorimotor integration center linking sensory threat information and escape. Adaptive changes in innate escape after learning could take a maladaptive turn after severe stress. Posttraumatic stress disorder (PTSD) is associated with long-term maladaptive changes after exposure to traumatic events, related to enhanced threat detection and reaction. Such long-term modifications are thought to involve the medial prefrontal cortex (mPFC), which is implicated in integrating learned emotional values into decisions that drive actions and behaviors. Here, in a series of experiments, we establish the crucial physiological role of specific mPFC neurons, exerting influence on the SC both directly and indirectly through the basal ganglia, in threat detection and reaction after adversity.

## Introduction

Innate defensive responses are crucial for survival, allowing split-second reactions to a potential threat^1, 2^. While primarily instinctive, threat responses are flexible and adapt to risk assessment by the organism, based on factors such as the perceived threat value of an agent in the environment, environmental conditions, internal states and prior knowledge of the organism^3, 4^. Modifying threat response is generally an adaptive process^5-7^, preventing unnecessary costs from the organism (in energy spent, feeding opportunities lost etc.)^8^. However, threat response flexibility could take a maladaptive turn after severe stress. Posttraumatic stress disorder (PTSD) is associated with long lasting changes after an exposure to a traumatic event, including changes in response to sensory information expressed in enhanced threat detection and reaction, and in reduced arousal thresholds and filter mechanisms^9-12^. As in human patients, major, long-lasting behavioral changes are evident in animals after a single exposure to a severe stressor^13-16^.

Escape behavior is a classical innate defensive action which, although required to be fast and almost reflex-like, includes intricate processing and computations to minimize reaction time while maximizing success or adaptivity by considering as much information as possible^3^. As such, in many species it is carried out by subcortical structures, directly receiving sensory information (bypassing sensory cortices) and engaging action, which allows for rapid and intense responses^17^. An important neural hub in managing escape is the superior colliculus (SC)^18-20^, a subcortical structure which receives direct retinal input on its superficial layers and generates premotor activity in the deeper layers^21, 22^, making it a subcortical sensorimotor integration center. Integration in the SC was shown to be crucial for linking sensory threat information and escape^23^. Importantly, adaptation of escape after learning was suggested to also occur in SC neurons^5, 18^, yet the source of this neural adaptation remains unknown. As escape response is carried out through rapid semi-instinctive systems, such long lasting modification may involve changes in higher control systems regulating threat responses.

The medial prefrontal cortex (mPFC) is implicated as a part of the prefrontal cortex which integrates previously learned emotional values into decisions and exerting adaptive control over different actions and behaviors^24-26^ depending on the specific subcortical circuit receiving mPFC input^27^. For example, activating mPFC projections to the dPAG, a specific relay station in the defensive behavior pathway, increased specific stress related actions, such as digging and marble burying^28^. The SC is also regulated by the mPFC, both directly^29, 30^ and indirectly through the basal-ganglia^31, 32^, where it is considered the cortical hub of the associative loop^33^, and involved in goal-directed action control^34^. Interestingly, it was recently demonstrated that this two-way cortical control of the SC could be traced to the same neurons in another frontal cortex region - the somatic sensorimotor area, which sends projections both directly to the SC and to the dorsal striatum which affects the SC via the cortico-basal ganglia-pathway^29^.

To find the mechanism by which escape threat response is modified after a significant aversive experience we recorded SC neurons of freely behaving mice confronted with erratically moving robo-beetle^35^. Behavior was recorded before and at two time points after a significant, unexpected and inescapable foot shock - a procedure known to elicit anxiety related long term changes in behavior^13, 15^. Following the aversive experience, we found long term changes in innate escape behavior - an increase in amount and intensity and an early onset, all correspond to changes in threshold of threat perception. Photo-tagging of incoming projections into the SC indicated that the changes were induced by mPFC neurons which project to both dorsomedial striatum (DMS) and directly to the SC. Examining general inputs into the SC either through general mPFC neural population or through all basal ganglia channeled input via the substantia nigra pars reticulata (SNr) seemed to dilute the change. Chemogenetically inhibiting mPFC neurons projecting to DMS during the aversive experience prevented the long-term change in escape threshold, while optogenetically activating their SC projections during BMT modified the threshold according to the distance-dependent stimulation. Our results point to specific top-down influence through a specialized circuit as necessary and sufficient in mediating enhanced threat detection after a significant aversive experience.

## Results

### Significant, unexpected, and inescapable foot shock induces long term changes in behavior and specifically in escape behavior

We first set out to test whether a severe, distressful experience (see ‘shock phase’ in the methods section) can alter the animal’s behavioral outcomes in context of a previously non-threatening stimulus and environment. To this aim we adapted the Beetle Mania Task (BMT)^35^ to our needs and changed it into a within subject design (**Figure 1a**). This allowed us to examine behavioral changes of the same animal in a semi-ethological environment, where the mouse encounters a smaller erratically moving robo-beetle. This moving, interacting stimulus has no preset threat value, nor does it harm the mouse in any way during interaction. Hence, it does not become a threat by interaction alone (**Figure 1b**). The mouse encounters the robo-beetle on three occasions – the first is prior to the shock, and then at two time points thereafter (**Figure 1a**). All interactions were recorded using a camera, set up above the arena, and locations and poses of both, mouse and robo-beetle, were identified and estimated using DeepLabCut^36, 37^. An escape was defined as an abrupt change in the mouse’s velocity, increasing the distance between itself and the approaching beetle. To characterize the changes in escape behavior other than the overall count of occurrences, we also measured the vectors of velocity and distance between mouse and beetle around escape onsets (**Figure 1c**). As a more general assessment for anxiety, we chose to measure the relative time the animal spent in the arena center (central 40% of arena range). Compared to the pre-shock baseline, we found that exposure to a significant aversive event resulted in increased numbers of escape responses as measured 7 days thereafter; a change which was maintained also 21 days post-shock (**Figure 1d;** a mixed model repeated measures ANOVA showed a significant main effect of time; F(2,44)=12.80, p<0.001; post-hoc test revealed a significant difference from the pre-shock baseline at the 7th day; p<0.01 and the 21st day p<0.001 from shock). Not only were there more escape events when measured 7 and 21 days after the shock, but we found that mice maintained a larger safety distance from the beetle as they detected it as a threat earlier and escape onsets occurred at a farther distance from the approaching beetle. (**Figure 1e;** a mixed model repeated measures ANOVA showed main effect of time; F(2,44)= 4.304, p<0.05; post-hoc tests revealed both 7^th^ and 21^st^ post-shock days differed from baseline; both p<0.05). The escapes events themselves where faster on average as they reached a higher peak of velocity during the escape both 7 and 21 days after the shock, as compared to baseline. (**Figure 1f;** a mixed model repeated measures ANOVA showed main effect of time; F(2,43)=5.572, p<0.01; post-hoc test revealed a significant difference from the pre-shock baseline at the 7th day; p<0.05; and the 21st day; p<0.01; post-shock). Those changes in escape behavior occurred after the shock administration and were maintained for weeks. Another maintained change was general anxiety as mice spent significantly less time in the central 40% of the arena on both the 7^th^ and 21^st^ day after the shock, as compared to baseline. (**Figure 1g;** a mixed model repeated measures ANOVA showed main effect of time; F(2,45)= 5.260, p<0.01; post-hoc tests revealed both 7^th^ and 21^st^ post-shock days differed from baseline; both p<0.05). All those changes could not be explained as resulting from the mere encounters with the beetle, as no such effects were observed in mice undergoing the same paradigm but without shock administration. (**Figure 1 h-k;** No significant effects of time in a repeated measures ANOVA of neither count, distance’, peak velocity or time spent in the central area all p= ns;).

**Figure 1.**
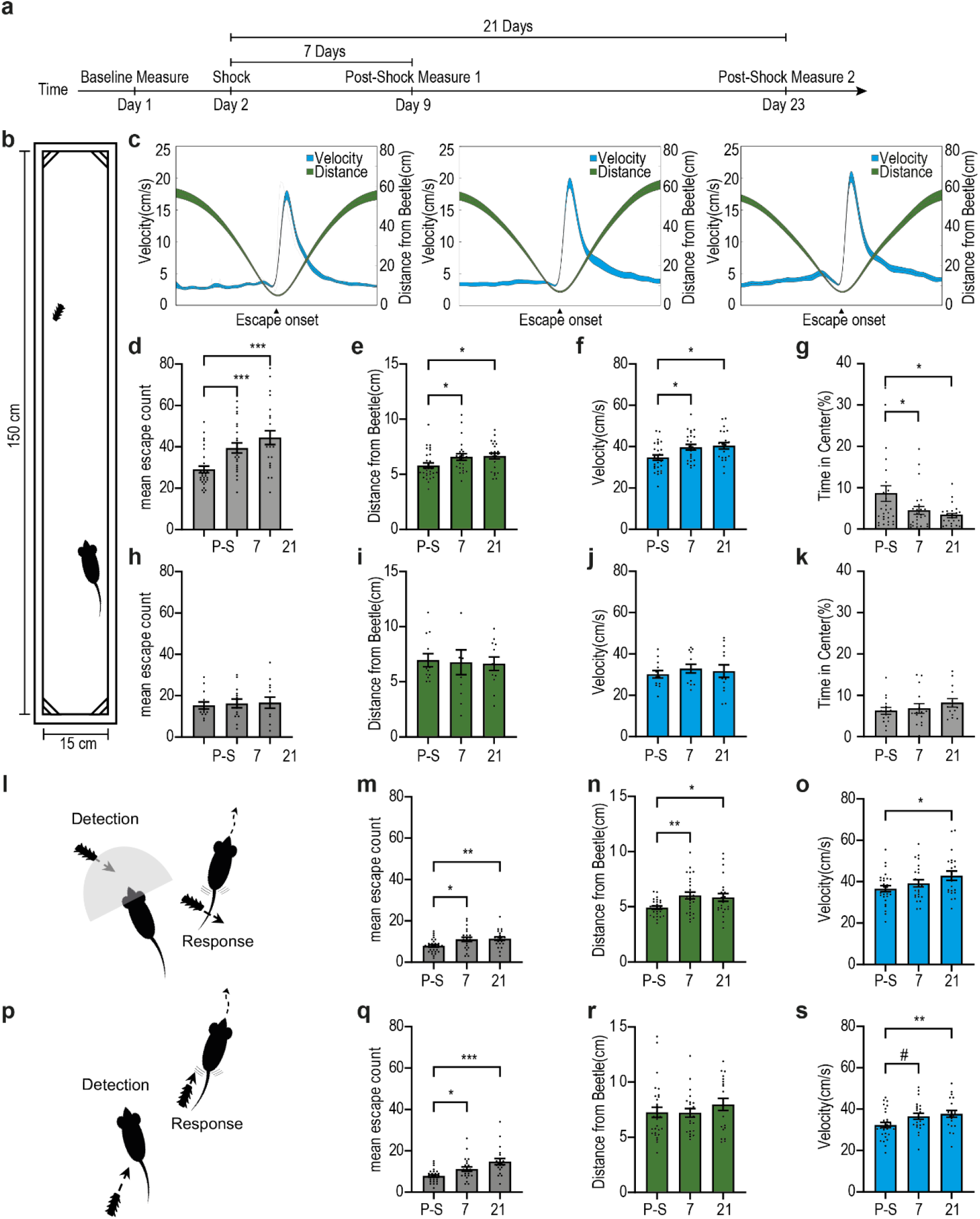
Exposure to severe adversity results in lasting changes in behavior. **(a)** behavioral paradigm and time schedule. **(b)** schematic depiction of the testing arena with the animal (bottom) and the beetle (top) to scale. **(c)** peri-event histograms depicting the animal’s velocity in cm/s (blue, left Y-axis) and the distance between mouse and beetle in cm (green, right Y-axis) around escape onsets, assessed pre-shock, 7 days and 21 days following footshock, left to right respectively. Each line represents the overall average of the corresponding measure with shaded ±SEM. **(d-g)** Behavioral measures in animals that received the foot-shock. **(d)** Significant Increase in number of escapes from beetle **(e)** Significant increase in the distance from beetle at escape onset (decrease in proximal tolerance to beetle) **(f)** Enhanced peak velocity of escapes. **(g)** A significant decrease in the time spent in the arena center. **(h-k)** The same behavioral measures in control animals (same schedule without footshock), no behavioral change was observed in general escape count **(h)**, distance at escape onset **(i)**, peak velocity of escapes **(j)** and in time spent in arena center **(k). (l)** An illustration depicting our definition of preventive escapes. The sketch displays a mouse and an approaching beetle. The gray half circle symbols the animal’s viewing direction and angle. An escape response following visual detection of the beetle by the animal was termed preventive escape (PE). **(m-o)** An increase in the number of PE from beetle **(m)**. A greater distance from beetle at PE onset. **(n)** Enhanced peak velocity of PE **(o)** following footshock. **(p)** An illustration depicting our definition of reactive escapes (RE) – the contrasting subtype for PE. The sketch displays a mouse with a beetle approaching from its rear, prompting the animal to escape once detected. **(q-s)** Similar to general escapes and PE, an increase in the number of RE **(q)** and enhanced peak velocity during RE onset **(r)** was observed post-shock and persistent over time. No change in distance at escape onset was observed **(s)**. In each plot, bars represent means of corresponding measures with error bars showing ±SEM. In peri-event histograms, each line is representing the overall average of the corresponding measure with shaded ±SEM. Asterisks indicate significant post-hoc comparisons (*, p<.05; **, p<.01; ***, p<.001). Pound signs indicate non-significant tends (#, .05<p<.1).

To isolate the effect of enhanced defensive distance, we further divided the escape events into escapes, initiated when the approaching beetle was in the line of sight of the mouse, termed - preventive escapes (**Figure 1l**) and escapes initiated when beetle was out of sight (usually by bumping into the mouse), termed - reactive escapes (**Figure 1p**). Some of the post-shock changes in escape characteristics occurred in both types of escapes. Such was the increase in the number of escape events occurring - a mixed model repeated measures ANOVA showed main effect of time for both preventive (**Figure 1m**; F(2,41) = 7.024, p<0.01; post-hoc test 7^th^ and 21^st^ day both p<0.01) and for reactive escapes (**Figure 1q** ; F(2,68)=11.13, p<0.001; post-hoc test 7^th^ day p<0.05 and 21^st^ day p<0.001). This was also true for the changes in peak velocity of escape, where repeated measure ANOVA revealed main effect of time for both preventive (**Figure 1o**; F(2,44)=3.252 = 7.024, p<0.01; post-hoc test 7^th^ and 21^st^ day both p<0.05) and for reactive escapes (**Figure 1s**; F(2,42)=4.497, p<0.05; post-hoc test 7^th^ day p=0.06 and 21^st^ day p<0.05).

However, changes in escape characteristics depending on visual detection, such as the safety distance, indeed occurred only in preventive escapes (**Figure 1n**; F(2,46)=4.660, p<0.05; post-hoc test 7^th^ and 21^st^ day both p<0.05). No such effect was found for reactive escapes. (**Figure 1r;** No significant effects of time in a repeated measures ANOVA ns).

### Superior colliculus (SC) neurons involved in escape initiation, maintain a longer safety distance long after experience with an inescapable foot shock

Searching for the neural changes underlying the early escape response within an enhanced threat detection distance, we implanted a custom made optrode (see methods section) in the SC of mice running the task (**Figure 2a-top**). We then recorded a total of 1395 units from the SC of 31 mice (see **Figure 2a-bottom** for the reconstruction of fiber locations). 80% (1115) of the recorded units significantly increased firing rate around events of any abrupt accelerations by the mouse and were termed “responsive to acceleration”, out of which 77% (864 units) were responsive specifically during preventive escapes, and 54% (630 units) responded during reactive escapes (**Figure 2b**). A vast majority (87%) of acceletation-responsive units initiated neural response prior to the acceleration itself (**Figure 2c-d**), indicating an involvement of those units in escape initiation as was previously suggested^38^. This distribution was solidly maintained when looking at the responses of recorded SC units at all three time-points of the task and even shifted to further precede acceleration on the 21^st^ day after the shock. (**Figure 2e left;** A Kruskal-Wallis test for assessing the differences in the distributions of neural responses initiation times relative to escape onset revealed a significant main effect of time H(2) = 49.65, p <0.001; post-hoc test 21^st^ day p<0.001). While the distance to the beetle at the time of neural response initiation preceding acceleration in acceleration-responsive units was maintained around the same distance (**Figure 2e right;** No significant main effect of time was found F(2,929)=0.0162, ns). To further study if this shift is related to escape behavior, we looked specifically at escapes through the prism of maintaining a defensive distance and separated to preventive and reactive escapes. In preventive escapes we found that the distribution moves further afore acceleration, slightly on the 7^th^ and significantly on the 21^st^ day post shock (**Figure 2f left;** Kruskal-Wallis test revealed a significant main effect of time H(2) = 33.05, p <0.001; post-hoc test 21^st^ day p<0.001), as well as the distance from the robo-beetle at the time of neural response onset (**Figure 2f right;** one way ANOVA revealed a significant main effect of time F(2,657)=4.408, p<0.05; post-hoc test 21^st^ day p<0.05). This however was not the case in responsive escapes where safety distance remained constant. (**Figure 2i-j;** H(2)=4.464, ns; as well as F(2,422)=1.351, ns).

**Figure 2.**
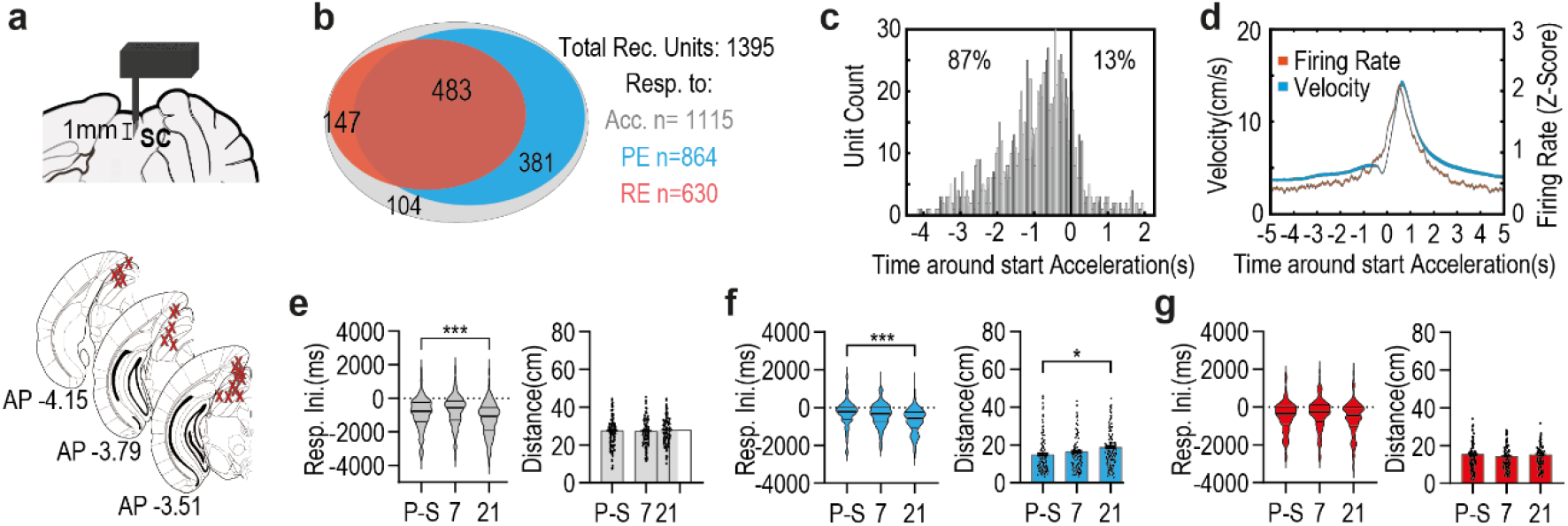
Altered safety distance following footshock as expressed in the neural activity of escape related SC neurons. **(a)** schematic depiction of the optrode design and placement (top). The optrode, was placed unilaterally in the SC. The bottom illustration depicts anatomical target schemes for all animals included in the experiment. X’s mark histologically confirmed most ventral position of the implant. **(b)** Diagram representing the distribution of acceleration responsive units with respect to whether they respond to PE, RE, both or neither. Numbers on each area of the Venn diagram correspond to unit count. **(c)** Histogram of neural response initiation time (s) in relation to acceleration onset (at time 0) for all acceleration-responsive units recorded in the SC. 87% of this population started responding before acceleration onset, while 13% started thereafter. **(d)** averaged neural response depicted in red (z-score of firing rate, right Y-axis) and averaged velocity depicted in blue (cm/s, left Y-axis) around acceleration onset (at time 0, X-axis). each line represents the overall average of the corresponding measure with shaded ±SEM. **(e)** In pooled acceleration-responsive units, neural response initiation times (ms) relative to acceleration onset undergoes a shift afore in distribution 21 days following footshock (left), yet no change in distance (cm) from the beetle at response initiation time is observed (right). **(f)** In PE responsive units, spike initiation times (ms) relative to escape onset undergoes a shift afore in distribution (left), as well as an increase in distance (cm) from beetle corresponding to response initiation time 21 days following footshock (right). **(g)** in RE responsive units, no change in time (left) or distance (right) at neural response initiation time could be observed at any timepoint following footshock. In each plot, bars represent means of corresponding measures with error bars showing ±SEM. In violin graphs, the bold midlines represent medians with IQR in regular lines above and below them. Asterisks indicate significant comparisons (*, p<.05; ***, p<.001).

While the persistent correlation between escape onsets and the preceding neural response suggests a direct involvement of SC neurons in the initiation and/or upkeep of escape behavior, the shift in the neural activity following shock, suggests upstream modulation of the SC involvement.

### Post-shock changes in preventive escape related activity does not map onto SC neurons receiving direct SNr Projections

To identify the neural inputs supporting the changes in the response timing of escape-SC neurons following the fearful event, we expressed an opsin in SC afferents while photo-tagging the neural response in the SC. We started by examining the output from the basal ganglia to the SC through the inhibitory projections from the SNr. A viral vector driving the expression of an opsin (pAAV-Syn-Chronos-GFP) was injected to the SNr and the optrode was implanted in the SC of mice running the task (**Figure 3a**). Using the optrode we recorded the units from the SC of 15 mice (see **Figure 3b** for reconstruction of the location of the fibers - left; and example microscope slice showing optrode damage+ opsin expression - right). SNr expression was also validated (**Figure 3c** for reconstruction of the opsin expression in the SNr - left; and example microscope slice – center and right). As shown in **Figure 3d** Out of 630 recorded units, 23% (143 units) were found responsive to SNr. 51% (73 units) of SNr responsive units were responsive specifically to PE (**Figure 3e)** and 41% (59 units) were responsive specifically to RE. The distribution of responses initiation was solidly maintained prior to acceleration at all three time-points of the task without a difference between them. (**Figure 3f left;** medians of - 340, -240 and -340 ms respectively with non-significant main effect of Kruskal-Wallis test H(2) =0.909; ns). No difference was also found in the distance to the beetle at neural response initiation preceding acceleration (**Figure 3f right;** No significant main effect of time was found F(2,50)=0.247, ns). The finding that the vast (73%) majority of SNr responsive SC units initiate response prior to PE suggests involvement of the SNr in escape initiation through its direct connection to the SC. As the temporal and spatial nature of this response stay stable following shock phase, this synapse does not seem to play a unique role in the behavioral and neural activity changes.

**Figure 3.**
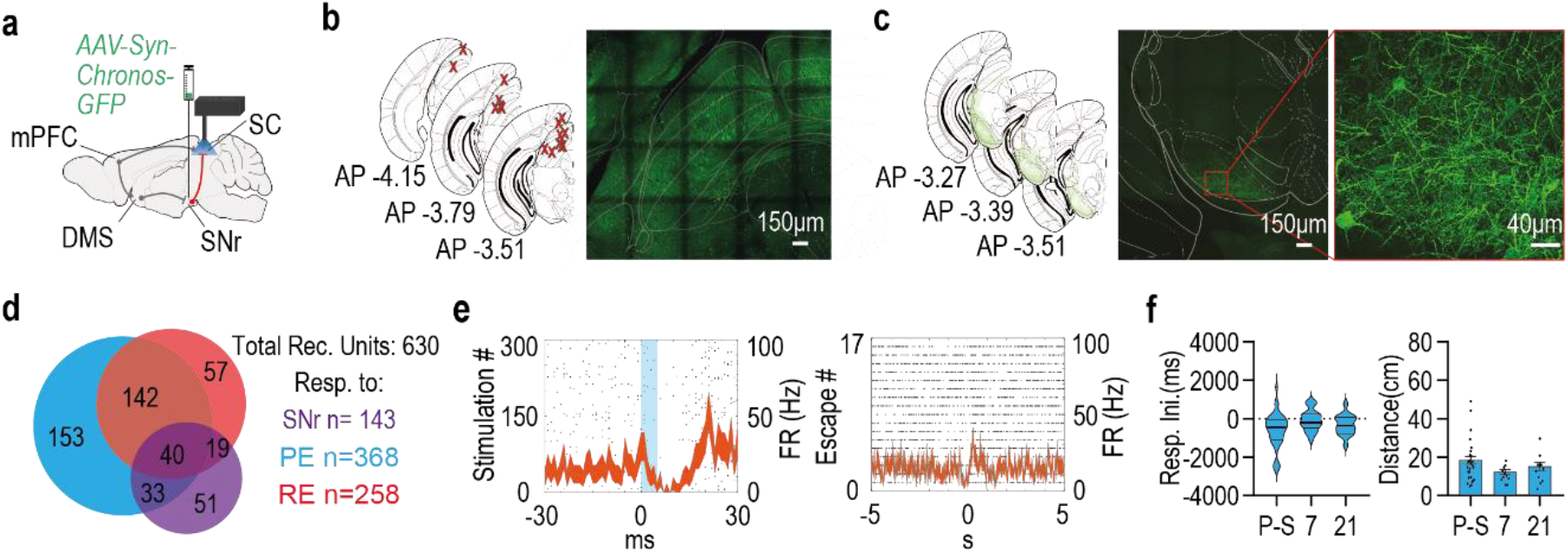
Altered safety distance following footshock is not expressed in the neural activity of specific SNr responsive escape related SC neurons. **(a)** schematic depiction of optrode location (SC) and viral injection site (SNr). **(b)** The left illustration depicts anatomical target schemes for all animals included in the experiment. X’s mark histologically confirmed implants positions. Right: representative coronal section showing the SC with traces of implant lesion and neural projections from the SNr. **(c)** The left illustration depicts overlaid, histologically confirmed SNr viral expression areas from all included animals. Center: representative coronal section showing viral expression in the SNr. Right: focusing on the SNr to see the projecting neurons. **(d)** Diagram representing the distribution of all recorded units in the experiment with respect to SNr responsiveness, as well as PE and RE related activity. Numbers on the Venn diagram correspond to unit count. **(e)** A raster plot overlaid with a PSTH showing a typical response of SNr responsive SC neuron to light stimulation (left) and to PE instances (right), each line represents the overall average of the corresponding measure with shaded ±SEM. **(f)** Response initiation times of PE related SNr responsive SC units almost exclusively precede PE onsets, but the distribution of response initiation times (ms) relative to PE onsets remained unchanged following footshock (left). No significant change was observed in distance (cm) from the beetle at the time of response initiation time of those neurons (right). In violin graphs, the bold midlines represent medians with IQR in regular lines above and below them, bars represent means of corresponding measures with error bars showing ±SEM.

### Post-shock changes in preventive escape related activity tracks down to SC neurons receiving direct and indirect input from a specific cohort of dorsomedial striatum (DMS) projecting medial prefrontal cortex (mPFC) neurons

To find out whether it is the prefrontal neural input which induces the changes in the response timing of escape-SC neurons following the fearful event, we tested two pathways by which the same mPFC neurons might exert influence over the SC, the first is through the basal-ganglia via the mPFC-DMS vast projections, the second is directly via mPFC-SC projections. In order to identify these unique inputs, we first injected a retrograde viral vector driving the expression of the Cre into the DMS (AAVrg-Ef1a-mCherry-IRES-Cre) and then a viral vector driving a Cre dependent expression of an opsin (pAAV-Ef1a-DIO hCHR2(E123T/159C)-EYFP) was injected into the mPFC. This allowed conditional expression of the opsin only in neurons projecting to the DMS. We then implanted an optic fiber was implanted above the DMS to allow the activation of mPFC terminals in the DMS and an optrode in the SC allowing the activation of mPFC terminals in the SC (of mPFC neurons projecting to the DMS) and recording SC neural activity (**Figure 4a**). Out of the total 366 units recorded 166 responded to photo-tagging of the DMS, 144 were responsive to mPFC direct projections, both of those neuronal types were mostly reactive during PE (**See Figure 4b** for the exact proportions of each category of response and **Figure 4c-e** for expression and implanting maps in the different regions). While these are assumably the same mPFC units which project both to the DMS and the SC we wanted to separate their direct and indirect influence on the SC neurons. Activating mPFC terminals in the DMS results in SC neural responses via the basal ganglia but might also result in antidromic activation which might also lead to SC response via the direct input of the same neuron. Likewise activating direct mPFC terminals in the SC could create an indirect SC response either via interneurons or antidromic activation. We therefore measured the response latency of SC neurons to light activation either above the DMS or directly above the mPFC and divided according to the type of response (excitation/inhibition). We found a short latency corresponding to one direct synapse only for SC neurons excited by direct mPFC projection activation^39, 40^, while neurons inhibited by direct mPFC projection activation, and both types of responses occurring in SC neurons by activation of mPFC terminals in the DMS, showed a significantly longer latencies (**Figure 4f**). A two-way ANOVA looking at the effect of response type (excitatory/inhibitory) and unit type (mPFC responsive/DMS responsive) on response latency after light stimulation found a significant interaction between the effects of response type and unit type was found (F(1, 346)=20.32, p <0.001) with multiple comparisons tests revealing a significantly shorter response latency in excitatory mPFC-responsive units compared to inhibitory mPFC-responsive units and all DMS responsive units (all p<0.001). We therefore considered only short latency (<10ms) SC neurons with excitatory responsive to direct SC light stimulation as direct mPFC responsive neurons and only SC neurons with long latency (>10ms) response to light stimulation of mPFC terminals in the DMS, as DMS responsive neurons. Example responses of DMS responsive and mPFC responsive units could be seen in **Figures 4g** and **4i** respectively. Interestingly, when isolating the SC neurons responsive to those DMS projecting mPFC neurons, either through the direct mPFC-SC synapse or through the DMS, we found that those neurons changed their response both around PE onset and relative to the distance from the beetle. So that the temporal distributions of DMS-responsive units around escape onsets changed from before to after the shock phase, as more units initiated their response earlier relative to PE onset (pre-shock: 66%, post 7 days: 92% and post 21 days: 89% - **figure 4h top**), indicating a quantitative shift in involvement in PE initiation (A significant main effect to time in Kruskal-Wallis test (H(2) =22.74, p <0.001) with post-hoc comparisons finding a significant difference of 7 and 21 days post shock from baseline both p<0.001). Moreover, DMS-responsive SC units also changed the time of response initiation from before to after the shock phase in relation to the distance from the beetle. After the shock phase response was initiated at greater distance from the beetle, reflecting the shift in the safety distance which was evident in the mice behavior (**figure 4h bottom)**. ANOVA revealed a significant effect of time (F(2,104)=5.478, p<0.01) with post-hoc comparisons finding a significant difference of 7 and 21 days post shock from baseline; p<0.01 and p<0.05 respectively).

**Figure 4.**
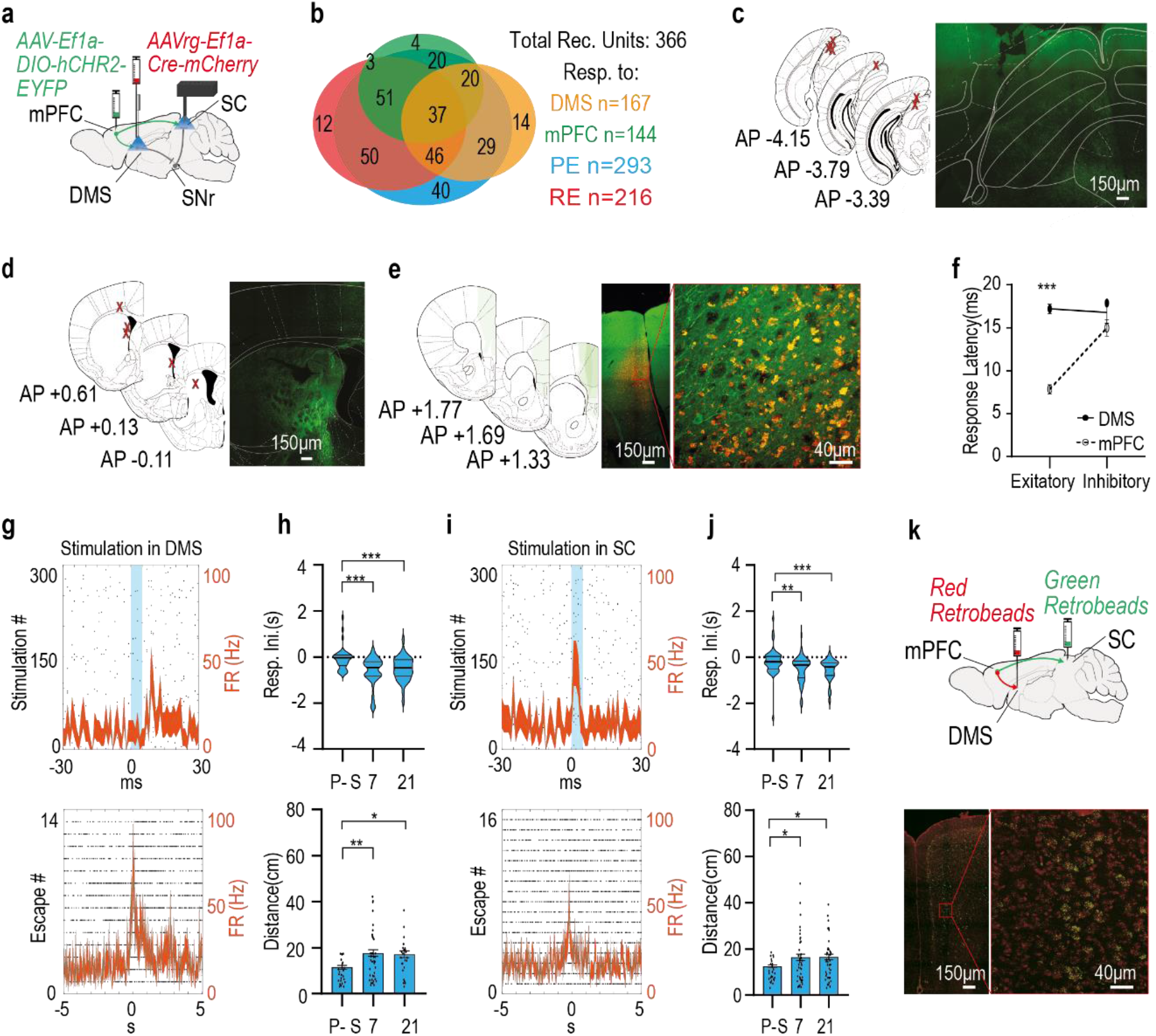
SC neurons affected directly or indirectly (through the DMS) by mPFC input, change the neural response to support behavioral change after the shock phase. **(a)** schematic depiction of optrode placement (SC), optic fiber position (DMS), and viral injection sites (mPFC and DMS). **(b)** A Venn diagram representing the distribution of all recorded units in the experiment concerning DMS and mPFC responsiveness, as well as PE and RE responsiveness. Numbers on the diagram correspond to unit count. **(c)** Left: anatomical target schemes showing the SC. X’s mark histologically confirmed optrode positions from all animals included in the experiment. Right: representative coronal section showing the SC with optrode lesion. **(d)** Left: histologically confirmed optic fiber placements at the DMS, marked by X’s, from all included animals. Right: representative coronal section showing the DMS with fiber lesion and neural projections from the mPFC in green fluorescence. **(e)** The left illustration depicts overlaid, histologically confirmed viral expression areas at the mPFC from all included animals. Right: representative coronal section showing viral expression and the co expression of red and greed indicating neurons of the mPFC projecting to the DMS. **(f)** Latency of SC neural response to photo-tagging, divided according to the type of response (excitation/inhibition). The full line depicts phototagging of SC neurons responsive to activation of mPFC terminals in the DMS, dashed line are units responsive to direct activation of mPFC terminals in the SC. Only units excited by mPFC direct projection activation show a short latency typical to a direct synapse. **(g)** A raster plot overlaid with a PSTH showing a typical response of the same SC neuron to light stimulation of mPFC terminals in the DMS (top) and to PE instances (bottom). **(h)** The distribution of response initiation times (ms) relative to PE onset in SC units responsive to mPFC terminals activation within the DMS, undergoes a sharp, significant shift afore following footshock (top) as the distance (cm) from the beetle (bottom) at response initiation by the same units increases. **(i)** Typical response of the same SC neuron to light stimulation of mPFC terminals in the SC (top) and to PE instances (bottom). **(j)** The distribution of response initiation times (ms) relative to PE onset in mPFC-responsive SC units, undergoes a sharp, significant shift afore following footshock (left) as the distance (cm) from the beetle (right) at response initiation by the same units increases. Both effects lasted for at least 21 days post footshock. **(k)** Top: A schematic depiction of Retrobeads injection sites at the DMS (red) and the SC (green). (g) Representative coronal section showing the mPFC with co-labeled neurons with red and green beads. Indicating that these neurons project to both the DMS and the SC. In each plot, bars represent means of corresponding measures with error bars showing ±SEM. In violin graphs, the bold midlines represent medians with IQR in regular lines above and below them. In peri-event time histograms (PSTH), each line represents the overall average of the corresponding measure with shaded ±SEM. Asterisks indicate significant post-hoc comparisons (*, p<.05; **, p<.01; ***, p<.001).

The same differences in the same direction were found in SC units responding to direct mPFC terminal activation of the same mPFC to DMS projecting neurons. We found a shift in the number of neurons initiating their response earlier relative to PE onset (**figure 4j top** A significant main effect to time in Kruskal-Wallis test (H(2)=25.44, p <0.001) with post-hoc comparisons finding a significant difference of 7 and 21 days post shock from baseline both p<0.001). We also found their neural response to be initiated at a greater distance from the beetle after the shock, reflecting the shift in the safety distance which was evident in the mice behavior (**figure 4j bottom)**. One-way ANOVA revealed a significant effect of time (F(2,95)=4.126, p<0.05) with post-hoc comparisons finding a significant difference of 7 and 21 days post shock from baseline; both p<0.05). To confirm the presence of DMS projecting mPFC neurons which are branching also to the SC, 2 animals were injected with green fluorescent microbeads (<u>Retrobeads</u>, Lumafluor) in the SC and red Retrobeads in the DMS (**Figure 4k top**). The abundant number of neurons in the mPFC, labelled with both red and green, provides evidence for the frequency of such bifurcating mPFC neurons. (**Figure 4k bottom**). The change in the PE responses of mPFC and DMS responsive SC neurons was almost identical. Given that the mPFC neurons projecting to both regions are all identified as DMS projecting, it appears that these neurons have a unique contribution to the change in the safety distance maintained prior to escape. This idea is further supported by the results of testing the SC response to a general population of mPFC projection cells without restriction to DMS projection mPFC cells. Mice were injected with viral vectors driving the expression of light-sensitive proteins (AAV5-CamkIIa-hCHR2-EYFP) in projection neurons of the mPFC and implanted with an optrode at the ipsilateral SC (**Supp. figure 1a, d-e**). Unlike in the experiment described before, the opsin expression in this case was not Cre-dependent. Thus, light stimulation above the SC evoked direct mPFC afferents, regardless of whether they also project to the DMS (or any other area) or not. Light-responsive units were termed ‘non-specific mPFC-responsive’ units (**Supp. figure 1b**). We found that non-specific mPFC-responsive SC units have a dimmed effect both on the distribution of response and on the distance from the beetle at PE onset (**Supp. figure 1c and d respectively**), with a significant effect only 21 days after the shock, much like the general effect on all PE responsive SC neurons (**Figure 2**). A significant main effect to time in Kruskal-Wallis test (H(2) =7.645, p <0.05) with post-hoc comparisons finding a significant difference only 21 days post shock from baseline both p<0.05). One-way ANOVA revealed a significant effect of time on distance (F(2,52)=7.42, p<0.01,) with post-hoc comparisons finding a non-significant difference trend 7 days post shock (p<0.08) and a significant difference 21 days post shock (p<0.001) from baseline. This again indicates a distinct contribution of the branched pathway in the altered, post shock phase neural response to PE.

### Silencing of DMS projecting mPFC neurons during shock phase prevents the enhancement of defensive distance

The physiological findings indicated that DMS projecting mPFC neurons play an important role in controlling the enhancement of defensive distance kept by SC neurons involved in escape behavior. To test whether indeed those neurons are necessary to this process we used HM4D DREADDs (designer receptor exclusively activated by designer drugs) to selectively silence DMS projecting mPFC neurons during shock phase. Mice were injected with a retrograde viral vector driving the expression of the Cre bilaterally into the DMS (AAV-retro-hEF1a-iCre) and then a viral vector driving a Cre dependent expression of either HM4D for the test group (AAV-CaMKIIa-dlox-hM4D-mCherry) or a fluorophore for control (AAV-CaMKIIa-dlox-mCherry) bilaterally to the mPFC (See **figure 5a** for injection scheme and **5c** for expression). The mice were put into the adapted BMT task and Clozapine-N-oxide (CNO) was administered 20 minutes’ prior the beginning of the shock phase in order to conditionally silence DMS projecting mPFC neurons during that phase (**figure 5b**). We found that mice not expressing HM4D show the expected increase in defensive distance 7 and 21 days following the aversive event. However, mice with selective silencing of DMS projecting mPFC neurons during the shock phase failed to show such an increase. Using a repeated measures mixed model ANOVA, we found a significant time X group interaction (F(2, 35) = 3.268, p<0.05). Post-hoc test revealed a significant increase in defensive distance between pre-shock to 7 days (p<0.01) and 21 days’ post shock (p<0.05) but not between any time point in the HM4D group (**figure 5d**). This suggests that DMS projecting mPFC neurons are required during the adversity in order to develop such an aversive experience dependant increase in defensive distance.

**Figure 5.**
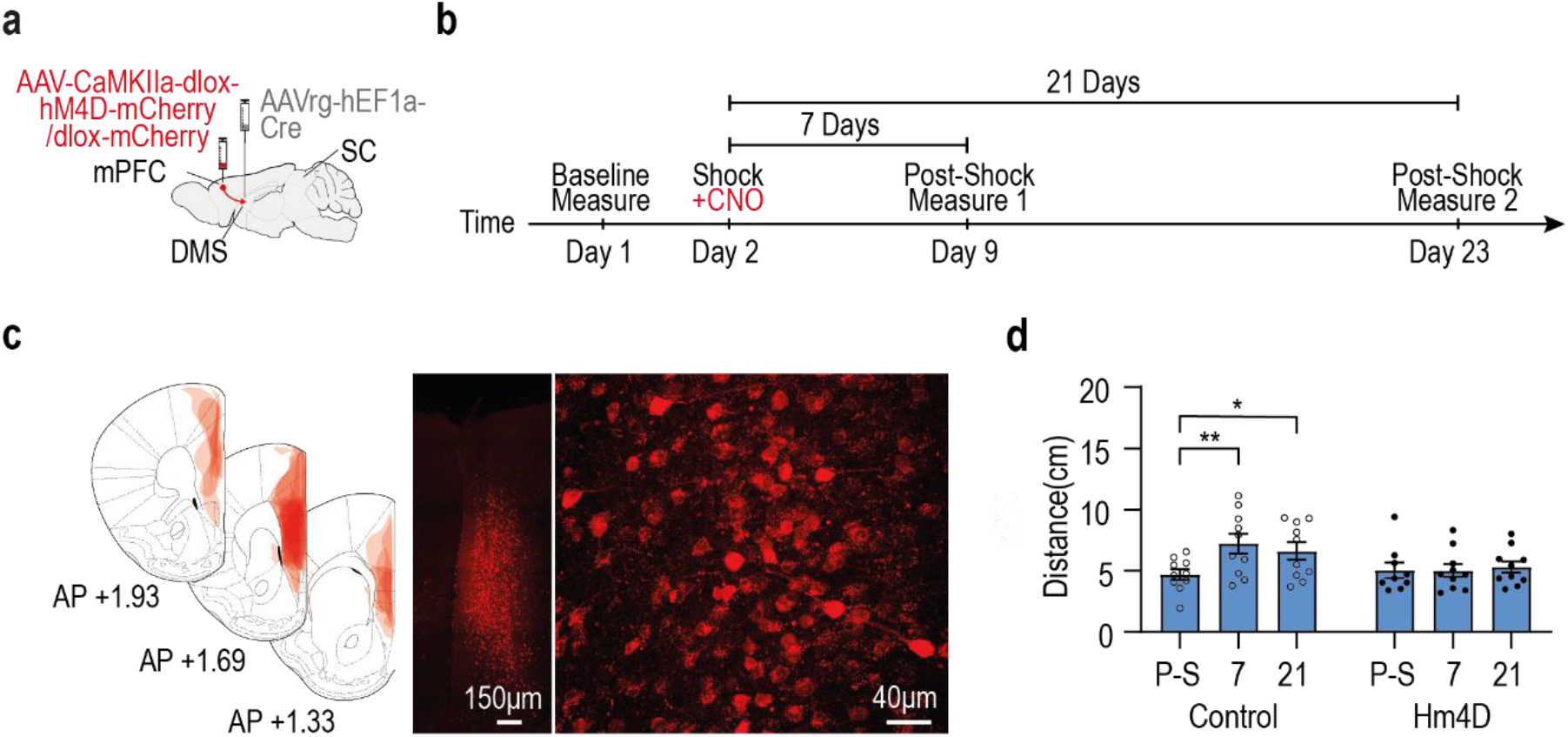
Silencing of DMS projecting mPFC neurons during shock phase prevents the enhancement of defensive distance. **(a)** Schematic depiction of viral injection sites (mPFC and DMS). **(b)** Experimental schedule. **(c)** The left illustration depicts overlaid, histologically confirmed viral expression areas at the mPFC from all included animals. Right: representative coronal section showing viral expression and neural cells in the mPFC. (d) Increased defensive distance following shock phase in control animals, but no difference following shock phase in Hm4D group. In each plot, bars represent means of corresponding measures with error bars showing ±SEM. Asterisks indicate significant post-hoc comparisons (*, p<0.05; **, p<0.01).

### Activating the SC terminals of DMS projecting mPFC neurons during the BMT controls defensive distance as a function of the distance from the approaching beetle

Finally, we tested whether those specific mPFC neurons which project to the DMS but also directly to the SC, are sufficient to change the defensive distance for escaping behavior, in naïve animals. Mice were injected with a retrograde viral vector driving the expression of the Cre bilaterally into the DMS (AAV-retro-hEF1a-iCre) and then a viral vector driving a Cre dependent expression of either an opsin for the test group (pAAV-Ef1a-DIO hChR2(E123T/T159C)-EYFP) or a fluorophore for control (pAAV-hSyn-EYFP) bilaterally to the mPFC. Optic fibers were implanted bilaterally above the SC. Thus, when stimulating, SC afferents from the branched mPFC neurons are selectively activated (See **figure 6a** for injection scheme and **6c-d** for fiber placement and expression in the SC and mPFC respectively).

**Figure 6.**
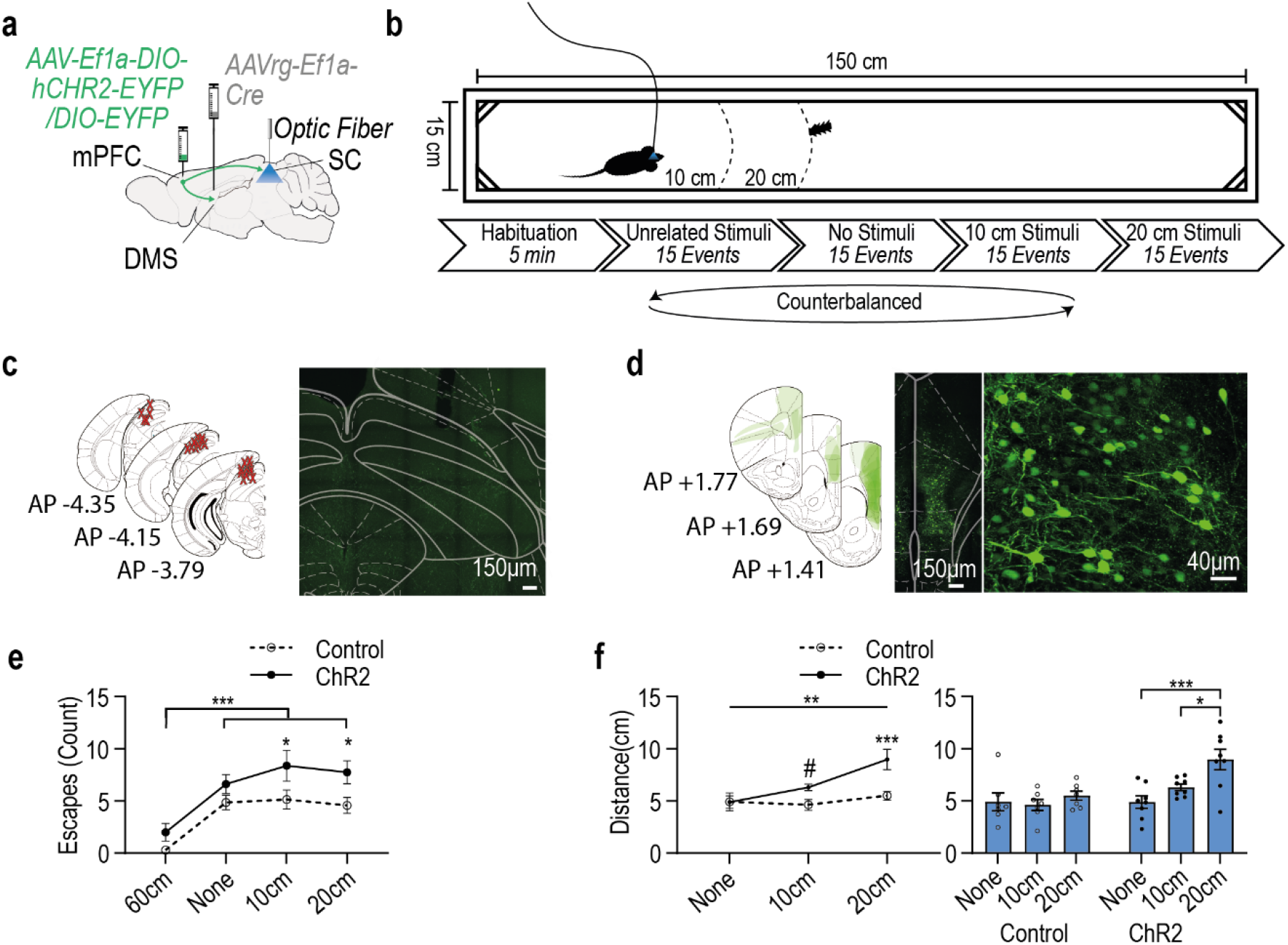
Activation of the terminals in the SC of DMS projecting mPFC neurons during a beetle’s approach affects defensive distance. **(a)** Schematic depiction of viral injection sites (mPFC and DMS) and fibers’ location. **(b)** Experimental setup and schedule. **(c)** Left: anatomical target schemes for all animals included in the experiment. X’s mark histologically confirmed optic fiber positions. Right: representative coronal section showing the SC with optic fiber lesion. **(d)** The left illustration depicts overlaid, histologically confirmed viral expression areas at the mPFC from all included animals. Right: representative coronal section showing viral expression and DMS projecting neurons in the mPFC. **(e)** Average (and SEM) of PE instances in each of the different stimulation phases devided to opsin expressing (ChR2) and fluorophore expressing (control) animals. (**f**) Average (and SEM) of the distance between the mouse and the beetle on PE initiation (defensive distance), in each of the different stimulation phases devided to opsin expressing (ChR2) and fluorophore expressing (control) animals.

The behavioral measurement was divided into 4 counterbalanced phases (following a short habituation). Each phase lasted until 15 mouse-beetle encounters occurred upon which stimulation was activated to a total of 15 light stimulations (1sec, 20Hz; corresponding to the average duration and frequency of the response to escape recorded in the SC). In each phase the stimulation was activated at a different encounter distance: 10cm, 20cm, 60cm (unrelated stimuli – this distance made sure that there was no real encounter during stimulation) and “none” (related to 15 beetle encounters without simulation-**figure 6b**).

To see whether the activation of the SC terminals of DMS projecting mPFC neurons is by itself sufficient to induce escape behavior, we compared the number of escape episodes (PE) of opsin and control animals occurring in each of the different phases. We indeed found that stimulation increased the number of escapes in the different phases, with the difference being focused on stimulation at 10cm and at 20cm phases (a repeated measures mixed model ANOVA found a significant main effect for opsin F(1, 13)=6.468, p<0.05; post – hoc comparison showing opsin is different than control only at 10cm and 20 cm phase, both p<0.05). This indicates that the stimulation increased the number of escapes only when the beetle stimulus was drawing near. We could also see the near absence of escapes at the 60 cm phase, indicating that the mere activation of those projections without the presence of the beetle is not sufficient to initiate escape behavior (a significant main effect for test phase F(3, 39)=22.09, p<0.001; with Tukey’s multiple comparison tests showing that animals at phase 60cm had significantly fewer PE instances compared to all other phases; all p<0.001) (**figure 6e**).

As there were merely any escapes at 60cm phase, measuring the distance during escape onset was irrelevant. To assess the effect of activation of the SC terminals of DMS projecting mPFC neurons on defensive distance we analysed the distance between the mouse and the beetle on escape initiation during the three other phases. We found that in general the opsin expressing animals kept an increased defensive distance as compared to control fluorophore animals (a repeated measures ANOVA found a significant main effect for group F(1, 13)=13.70, p<0.01). The increase in defensive distance was a function of the distance on which stimulation was performed. That is, the larger the distance was between the mice and the beetle when stimulation commenced the larger was the defensive distance kept before escape was initiated (a significant main effect for test phase was also found (F(2, 26)=6.07, p<0.01) with post-hoc comparisons showing that animals at phase 20cm kept significantly increased defensive distances compared to phase 10cm as well as to phase none (p<0.05 and p<0.01 respectively). Furthermore, a statistical trend in the interaction effect was also found F(2, 26)=3.15, p=0.06 in which post-hoc comparisons revealed that only in the opsin condition animals at phase 20cm kept significantly increased defensive distances compared to phase 10cm as well as to phase none (p<0.05 and p<0.001) respectably, while no differences were found within the control group. Additionally, it revealed increased defensive distances between opsin and control only at stimulation phases: at phase 20cm (p<0.001), as well as a statistical trend to increase at phase 10cm between test control animals (p=0.07) (**Figure 6f**).

These findings indicate that the DMS and SC branching mPFC neurons are sufficient to induce escape behavior in response to an approaching stimulus. Not only that, but they are also sufficient to set a defensive distance as a function of the timing of their activity.

## Discussion

The present study was designed to find the physiological basis of top down, long term changes in innate threat responding occurring after a significant aversive event, as reported in cases of PTSD^9, 10, 12^. Specifically, how does a traumatic event, such as a severe set of inescapable foot-shocks, creates long term changes in escape responses to perceived threat and the mechanism by which such changes are generated in the SC by mPFC influence.

Indeed, we found significant changes in innate escape behavior of mice long after the aftermath of the shock procedure, mice not only escaped more frequently from a neutral, previously non-threatening stimuli (the robo-beelte) but also escaped earlier, when the beetle was more distant, thus keeping an increased safety- or defensive distance compared to pre shock. This suggests an enhanced sensitivity to potential threats and a decreased threshold to initiate escape. The escapes themselves reached higher velocities, which could also indicate the heightened degree of perceived danger from the approaching object. While most effects were general changes to all escapes the effects of adversity on defensive distance was specific to preventive escapes (PE) - corresponding to instances where visual detection of the oncoming threat preceded the elicited escape (mouse facing the approaching beetle), and did not appear in reactive escapes (RE) - where the beetle approached the mouse from behind (and the mouse did not turn its head toward the beetle), therefore not establishing visual contact prior to escape. This suggested that the change in defensive distance indeed requires visual detection.

Therefore, it might not be surprising that physiological changes supporting the shift in threat detection were indeed found in the SC, a midbrain structure known to integrate direct visual information and promote responses to visual threats^4, 17-20^. Our findings support the involvement of this structure in innate escape as 80% of the units recorded in this area responded with increased neural firing rate to the mouse sudden acceleration onsets, while a vast majority thereof (87%) initiated response prior to acceleration onsets, suggesting SC involvement in the generation of this behavior. When looking at the exact initiation time of the neural responses of those neurons relative to escape onsets, we found that an increasing number of them shifted response initiation to an earlier time prior to escape initiation and to a further distance from the perceived threat long after the adversity. Changes in the physiology of the innate defensive system after a traumatic experience was already described both in animals and in humans^41, 42^. Yet the systems driving such a change are still uncharted.

When assessing the contribution of different structures connecting into the SC, to the observed effects, we found that the quantitative change in neural engagement following shock, as well as the temporal and proximal shift of SC escape neurons could be clearly attributed to a unique set of neurons in the mPFC. The mPFC is thought to play a role in behavioral control related to emotions^24-26, 43, 44^ and induce behavioral changes due to changes in the emotional saliency and valence of a stimulus^45^. It also connects to the SC through its vast projections into the basal-ganglia^31, 32^ where it is thought to be the affective propagator of goal-directed behavior^46, 47^. Indeed, when testing the input of the mPFC into this system, by looking at the input reaching the SC from mPFC neurons projecting into the input nucleus of the basal ganglia - the DMS, we identified that SC escape neurons responsive to this input shift their response to support early escaping behavior. We also found that the effect of those DMS projecting mPFC neurons reached the SC both through the basal ganglia but also directly and in both ways, it supported the shift in the defensive distance. This effect was specific to those DMS projecting mPFC neurons, as when testing the general effect of SC neurons to direct mPFC projections, the effect was significantly milder. Also, when testing all input reaching the SC escape neurons from the basal ganglia through the SNr we could not find the specific shift after the shock treatment. That is, most of SNr responsive SC neurons responded prior to escape but we could not identify the ones that shifted within this population.

The specificity of the effect of DMS projecting mPFC neurons on the same escape neurons in the SC which shifted their response after the shock treatment, indicated that those unique mPFC neurons play a significant role in the physiological basis of the behavioral shift. The literature has long been describing long term changes in mPFC plasticity after fear^48^, stress^49, 50^ and in relation to PTSD^51, 52^. To test whether changes in the specific DMS projecting mPFC neurons contribute to the long-term change in defensive distance we selectively silenced them during the footshocks phase using DREADDS. We found that this selective silencing eliminated the long-term increase in defensive distance. This indicated that the activity of DMS projecting mPFC neurons during the adversity is necessary for this behavioral change to occur.

The indirect connection of those neurons to the SC through the DMS and the basal ganglia, could establish a long-term change in the selection of escape action. However, considering the importance of prompt and rapid responses to threat, an indirect connection to the SC, could prove suboptimal for immediate threat response. Yet, through direct mPFC - SC connection by the same DMS projecting mPFC neurons immediate influence can also be exerted. Notably, we found a bifurcating pathway, originating at the mPFC and branching both to the DMS and directly to the SC. Both by our photo-tagging and electrophysiological recording, and by histological verification, using retrograde fluorescent tracing.

Activating the direct terminals in the SC of the DMS projecting mPFC neurons of naïve animals running the task, at the average frequency and duration measured in the response of escape related SC neurons, resulted in an increase in the number of escapes but only when the beetle was actually approaching. This indicates that this pathway induces escape in relation to a target and not merely a burst in running velocity. Interestingly the distance between the mouse and the robo-beetle in which the stimulation was activated affected the distance in which the escape was initiated in, indicating that the activation of the direct SC projections of the DMS projecting mPFC neurons was sufficient to activate early escapes.

Taken together the experiments outlined in this work show that changes occurring in the almost reflex-like escape innate defensive response, following a significant dire experience, map onto neurons in the SC. Specifically SC neurons receiving input from a unique population of mPFC neurons that execute their effect on behavior both via the action-selection pathway of the basal-ganglia, but also directly on the SC. Those mPFC neurons can initiate escape from a perceived threat, and acquire alterations during adversity, rendering them more prone to earlier escape initiation through their connection to the specialized neurons in the SC. What the different functions, or purposes of the direct and indirect pathways are, and whether other such pathways originating in the mPFC are serving different defensive actions, are questions for future research.

## Materials and methods

### Animals

In total, 65 male C57BL/6 Mice, (60 days old, ∼25g; Envigo, Jerusalem, Israel), were housed together (up to 5 per cage) at 22 ± 2°C under 12-hour reversed light/dark cycles. Mice were allowed water and laboratory rodent chow ad libitum. The experiments were approved by the University of Haifa Ethics and Animal Care Committee, and adequate measures were taken to minimize pain and discomfort.

### Behavioral Paradigm

#### Beetle Mania Task (BMT)

Our modified version of BMT is based on a paradigm established at the Wotjak’s Lab it the Max Planck institute for Psychiatry^35^. First, mice are individually placed into a dimly lit, custom made, white acrylic arena (H37, L150cm, W15cm) (**figure 1a-b**) for 5 min habituation. Next, a randomly moving, battery operated toy beetle (H1.8 cm, L4.5 cm, W1.5 cm, weight: 7.3 g, mean speed: 25 cm/s; https://www.hexbeetle.com/nano, Innovation First Labs, Inc., TX, USA) is introduced into the arena. For the next 10 minutes, the arbitrarily moving beetle creates accidental contacts with the mouse, allowing us to observe the animal’s responses and general behavior. In our experiments this basic paradigm was first conducted with naïve animals to establish baseline measures and was repeated 7 days and 21 days following shock phase (described below) to examine possible differences in the behaviors of interest. Each trial was video-recorded in 25fps, using an overhead-mounted camera. In experiments involving electrophysiological recordings, 10 more minutes for photo-tagging were added to the schedule (5 after habituation and 5 at the end of the trial). In the final experiment the schedule was slightly altered (**figure 6b** for further discussion see the results section).

#### Behavioral Measures

All measures regarding the animals’ behavior were detected and extracted using DeepLabCut2^36^ and a custom made code in Matlab (Version R2021a). The study focused primarily on aspects of escape behavior. An escape was defined as a sudden acceleration of the mouse, in reference to the approaching beetle (during all phases of acceleration). Later we made further distinction between escapes in which the mouse faces the approaching beetle, termed *Preventing Escape (PE)* and escapes in which the animal is approached from behind, termed *Reactive Escape (RE)*. Where online detection of mouse-beetle proximity was needed, Ethovision XT (Noldus) was used for this task. Several behavioral measures were extracted:

##### Number of escape occurrences

Total count of escape instances as described.

##### Defensive Distance

Distance between mouse and beetle at escape onset. Our measure for proximal tolerance before threat response.

##### Escape Velocity

Peak mouse velocity (cm/s) following escape onset.

##### Time in Arena Center

Expressed in percentage of the total trial time the animal spent in the inner 40% of the arena (in accordance with the rectangular open field. This was used as an estimation for general anxiety.

#### Stressor (Shock Phase)

To induce a non-associative, PTSD-like anxiety phenotype, we used the established dual shocks as a mouse model of PTSD ^13, 53, 54^. Mice were individually placed into a shock chamber (Med-Associates, St Albans, VT), which was scented with 70% EtOH. After 198s, two strong electric foot shocks (2s, 1.5 mA) were presented, spaced apart by 40s. Animals were returned to their home cage 60s after the last shock. Control animals were placed into the shock chamber for 300s, without shocks.

### Physiological Procedures

#### Stereotaxic surgery

All mice were anesthetized by injection of a Ketamine-Xylazine Solution (0.04ml/20g, i.p.) and secured in a stereotaxic frame (Kopf Instruments). Isoflurane (1%) was delivered by a low-flow anesthesia delivery system (Kent Scientific) to maintain a deep anesthetized state over the course of the surgery. Body temperature was maintained at 37°C by controlling their temperature with a thermal probe and an automatic controller system. Mice received an injection of analgesic agent Carpofen (0.01ml/20g, i.p.) and ophthalmic ointment (Duratears) was applied on the mice eyes to protect the cornea from drying during the surgery. Following surgery, mice received systemic antibiotics (Amoxy LA, 0.04ml/20g, i.p.) for 4 consequent days (1injection every 48h) including the day of the surgery and were allowed to recover for at least 1 week before initiation of behavioral experiments. Depending on the specific experiment, optrodes, optic fibers, viral vectors or fluorescent tracers (Retrobeads, Lumafluor) were injected/implanted.

#### Optrodes

For electrophysiological recording and optogenetic stimulation at the SC, a custom made optrode was used, comprising of 16 tungsten wires and an optic fiber (Thor Labs), attached to an 18-pin connector (Mill-Max, NY). Optrodes were grounded via a wire placed in the cerebellum. The optrode was positioned unilaterally in the SC (AP:-3.80mm, ML:±0.8mm, DV:-1.5mm) and secured to the skull using C&B Metabond (Parkell, Edgewood, NY) and dental cement.

#### Recordings

Extracellular waveform signals were recorded using Digital Lynx hardware and software (Neuralynx Inc; USA). The signal was filtered (600-6,000 Hz) and amplified using a head-stage amplifier. Neural data was sorted manually using Offline Sorter (Plexon; USA) and analyzed in Matlab (Version R2021a).

Response to light stimulation (Phototagging) was used to classify units responsive to specific inputs. 300 light stimulations (λ=470, 5ms wide pulses) were admitted through the optical fibers/optrodes at 0.5Hz rate, 150 before and 150 after behavioral paradigm. Paired-samples t-tests were calculated to compare baseline neural activity to response time windows. For photo tagging the baseline window was based on the 30 ms prior to the delivery each light stimulation and was compared to 1ms to10ms for an early response, and 10ms to 20ms for a late response. For behavior, the baseline window was defined as 5 to 2 seconds prior to behavioral event, and compared to the two seconds around the event in bins of 400ms. Units that had at least one significant comparison (p<0.05) within were classified as responsive.

#### Viral vectors and beads

Viral vectors were injected using a Nanofil Syringe mounted on a micro injection syringe pump and a (World Precision Instruments, Sarasota, FL). The injected volume was 600nl at rate of 100nl per minute. After each injection, the needle was left in place for an additional 1 minute per 100nl and then slowly withdrawn. The skin was then resealed and mice were allowed to recover for at least 6-8 weeks to allow full expression of opsins. On unilateral injections vectors were injected ipsilaterally to optrode at the SC implanted later.

##### List of vectors and beads

For the investigation of the SNr-SC connection, pAAV-Syn-Chronos-GFP^55^ (Provided by Edward Boyden, Addgene plasmid # 59170) was injected to the SNr (AP:-3.24mm, ML:±1.4mm, DV-4.5mm), unilaterally, ipsilaterally to the implanted optrode at the SC.

For the investigation of the mPFC-SC connection, AAV5-CamKIIa-hChR2 (E123T/T159C)-eYFP^56^ (Provided by Karl Deisseroth; UNC,NC) was injected to the mPFC (AP:1.70mm, ML:±0.3mm, DV-2.7mm), unilaterally.

For the investigation of the mPFC-evoked DMS-SC connection, two viral vectors were injected. Into the mPFC (AP:1.70mm, ML:±0.3mm, DV-2.7mm), pAAV-Ef1a-DIO-hCHR2(E123T/159C)-EYFP^56^ (provided by Karl Deisseroth, Addgene plasmid # 35509). The second vector, AAVrg-Ef1a-mCherry-IRES-Cre^57^ (provided by Karl Deisseroth, Addgene plasmid # 55632) was injected into the DMS (AP:+0.5mm, ML:±1.25 mm, DV-3mm). Both vectors were injected unilaterally.

For chemogenetic inhibition of DMS projecting mPFC neurons, two viral vectors were injected. Into the mPFC (AP:1.70mm, ML:±0.3mm, DV-2.7mm), a CRE-dependent vector, anterogradely driving the expression of hM4D receptors for Clozapine-N-oxide (CNO)-induced neuronal silencing, ssAAV5-mCaMKIIα-dlox-hM4D(Gi)_mCherry(rev)-dlox-WPRE-bGHp^58^ (constructed by the VVF, Addgene #50461) or alternatively, for control animals, ssAAV5-mCaMKIIα-dlox-mCherry(rev)-dlox-WPRE-bGHp (constructed by the VVF) was injected. A second vector, retrogradely driving the expression of Cre, ssAAV-retro/2-hEF1α-iCre-WPRE-bGHp (constructed by the VVF, Addgene #24593) was injected into the DMS (AP:+0.5mm, ML:±1.25 mm, DV-3mm). Both vectors were injected bilaterally.

For optogenetic activation of DMS and SC projecting mPFC-afferents at the SC, two viral vectors were injected. Into the mPFC (AP:1.70mm, ML:±0.3mm, DV-2.7mm), a CRE-dependent vector, anterogradely driving the expression of light-sensitive proteins, pAAV-Ef1a-DIO hCHR2(E123T/159C)-EYFP^59^ (provided by Karl Deisseroth, Addgene plasmid # 35509) or alternatively, for control animals, rAAV5/hSyn-Con/Foff--eYFP-WPRE^57^ (provided by Karl Deisseroth, Addgene plasmid # 55651) was injected. A second vector, retrogradely driving the expression of Cre, ssAAV-retro/2-hEF1α-iCre-WPRE-bGHp (constructed by the VVF, Addgene #24593) was injected into the DMS (AP:+0.5mm, ML:±1.25 mm, DV-3mm). Both vectors were injected bilaterally.

##### Retrobeads injections

For anatomical tracing of SC afferents and DMS afferents, Green (to the SC) and Red (to the DMS) fluorescent tracer microspheres (Retrobeads, Lumafluor) were injected into the SC (AP:-3.80mm, ML:±0.8mm, DV:-1.5mm) and into the DMS (AP:+0.5mm, ML:±1.25 mm, DV-3mm). The stock solution was diluted in pbs (1/4) and the volume per injection area was 600μL.

## Acknowledgements

We thank Carsten T. Wotjak for the discussion and ideas during the work on this project, Matthias Prigge and all Klavir lab members for discussions and critical reading of the manuscript. This work was supported in part by grants from the German Israeli Foundation (GIF I-270-421.10-2016), and the Israel Science Foundation (ISF 1495/17).

