## Supplementary material for "Prefrontal Control of Innate Escape Behavior – A Neural Mechanism of Enhanced Posttraumatic Threat Detection": Supp. FIgure1

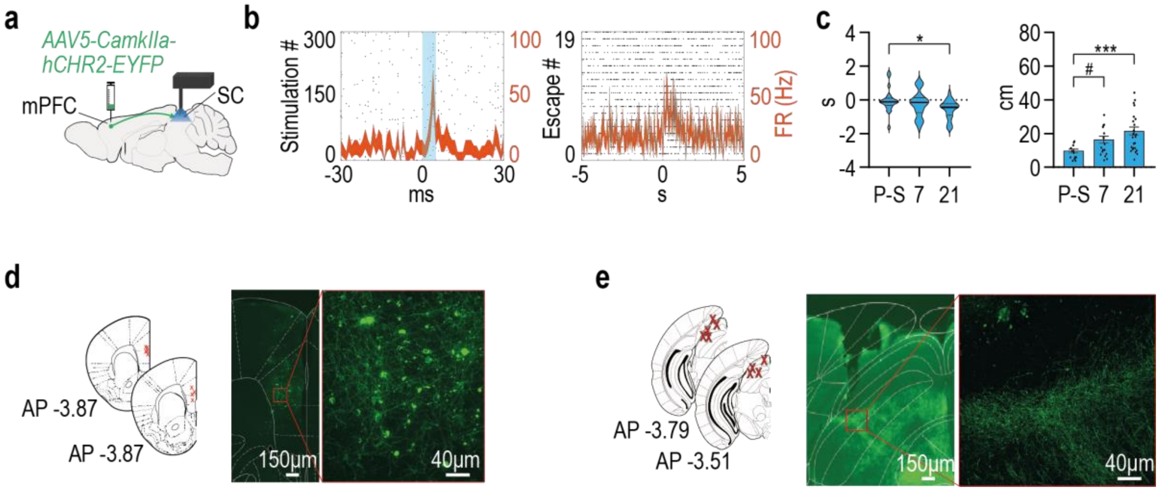


**Supplementary figure 1. SC neurons affected directly by general mPFC input, mildly change the neural response after the shock phase. (a)** Schematic depiction of optrode placement (SC) and viral injection site(mPFC). **(b)** A raster plot overlaid with a PSTH showing a typical response to light stimulation of mPFC terminals in the SC (left) and PE instances (right) of non-specific mPFC-responsive afferents at the SC. **(c)** In PE-responsive units, spike initiation times (ms) relative to escape onset seem to undergo a shift afore in distribution (left), as well as greater distance (cm) from beetle and spike initiation time (right) 21 days following footshock. **(d)** Histologically confirmed injection sites at the mPFC, marked by X's, from all included animals. Right: representative coronal section showing viral expression in mPFC projections cells. **(e)** Histologically confirmed optrode placement sites at the mPFC, marked by X's, from all included animals. Right: representative coronal section showing viral expression of mPFC projections in the SC. In each plot, bars represent means of corresponding measures with error bars showing ±SEM. In violin graphs, the bold midlines represent medians with IQR in regular lines above and below them. In peri-event histograms, each line is representing the overall average of the corresponding measure with shaded ±SEM. Asterisks indicate significant post-hoc comparisons (*, $p<0.05$; ***,$p<0.001$). Pound signs indicate non-significant trends (#, $p<0.1$ - $p>0.05$).
